# Upregulation of Calbindin in Adult Inhibitory Neurons Reactivates Critical Period Plasticity in Mouse Visual Cortex

**DOI:** 10.64898/2026.07.08.737337

**Authors:** Taylor Nakayama, Dario Figueroa-Velez, Whitney England, Emily Miyoshi, Jonathan Hasselmann, Robert C. Spitale, Vivek Swarup, Mathew Blurton-Jones, Jon T. Sack, Timothy J. Gardner, Sunil P. Gandhi

## Abstract

Critical periods are windows of peak learning performance where heightened synaptic plasticity enables fast, robust reorganization in juvenile brain circuits. Unlike adult plasticity which requires ongoing, persistent change in sensory experience, cortical representations can be rapidly changed during critical periods and have enduring effects. Transplantation of embryonic GABAergic inhibitory neurons has been shown to restore critical period plasticity to recipient neocortical circuits by triggering signalling changes within host inhibitory neurons. Here we transcriptionally profiled host inhibitory neurons in mouse primary visual cortex (V1) to detect gene expression changes during transplant-induced plasticity. Gene ontology enrichment analysis of differentially-expressed (DE) transcripts in host inhibitory neurons revealed synaptic plasticity- and inhibitory neuron development-related profiles. We assessed the protein expression of a top DE candidate, the calcium-binding protein Calbindin (Calb1), across transplant conditions and developmental stages. We found high Calb1 expression during the V1 critical period and during transplant-induced plasticity. To assess the functional significance of transplant-reactivated DE gene activity on visual cortical plasticity, we developed a set of AAVs to manipulate Calb1 expression specifically within inhibitory neurons. Using intrinsic signal imaging to measure ocular dominance plasticity, we found that Calb1 levels in V1 inhibitory neurons determine the extent of visual cortical plasticity. Our study provides evidence that induced expression of a critical period-stage gene in inhibitory neurons restores juvenile plasticity in targeted adult cortical circuits.

## Main Text

The maturation of inhibitory neurons triggers the opening of a brief critical period in which sensory experience can reorganize central circuits in the brain (*1*). After closure, the critical period can be reopened through the transplantation of embryonic inhibitory neurons into targeted brain regions (*2–3*). Despite evidence of robust and sustained functional recovery across diverse therapeutic applications (*4,5*), inhibitory neuron transplantation remains predominantly conceptualized as a cell-replacement strategy. The number of engrafted inhibitory neurons, however, does not correlate with extent of circuit reorganization (*6*), nor does their functional silencing prevent the induction of new plasticity (*7–8*). Recent findings from mouse visual cortex (V1) suggest that transplanted inhibitory neurons reactivate cortical plasticity by modifying their adult host counterparts to re-express an intracellular developmental program (*8*).

In this study, we analyzed RNA transcripts from adult host inhibitory neurons isolated from transplanted V1 during the period of transplant-reactivated cortical plasticity. From that transcriptomic profile, we validated the differential expression (DE) of candidate gene Calbindin (Calb1) during both the juvenile and transplant-reactivated period of cortical plasticity using immunohistochemistry. Given strong correlation between high Calb1 expression and cortical plasticity, we used viral-genetic targeting to specifically upregulate Calb1 in adult inhibitory neurons from non-transplanted V1. Together, the results indicate that Calb1 activity in inhibitory neurons plays a potent and direct role in regulating cortical plasticity.

### Isolation of host inhibitory neurons from transplanted adult V1

To genetically label host inhibitory neurons in recipient brains, we crossed transgenic mice expressing pan-inhibitory marker VGAT-Cre (Cre-recombinase under control of the vesicular GABA transporter promoter), with mice carrying Cre-dependent fluorescent reporter (ZsGreen) (Fig 1a). To distinguish mechanisms that govern transplant-reactivated cortical plasticity from endogenous juvenile plasticity, all engraftment was conducted in adult recipients (∼P120). VGAT-ZsGreen mice were affixed with custom-printed headplates centered over V1 on the right hemisphere. To account for anatomical variation between mice, V1 boundaries were physiologically determined prior to transplantation via intrinsic signal optical imaging (ISOI) (Fig 1). This functional imaging method leverages hemodynamic changes as a readout for neural activity and can be combined with visual stimuli to assess V1 responses (*9*). ISOI-generated retinotopic maps of binocular V1 were then superimposed onto images of surface vasculature to determine optimal injection sites.

**Fig. 1.**
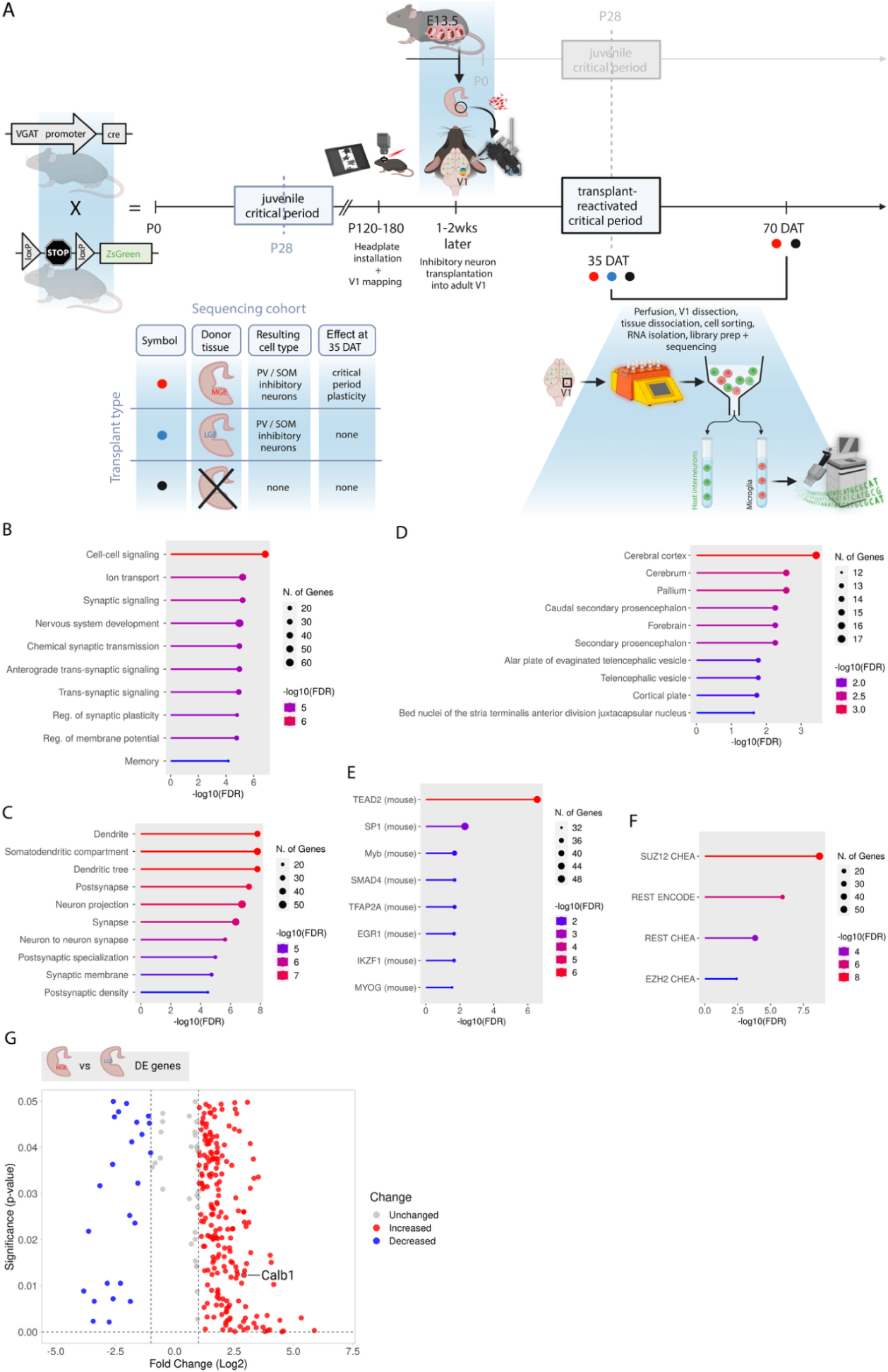
Comparison of host inhibitory neuron gene expression profiles from MGE vs. LGE transplanted adult visual cortex. **(A)** Schematic of experimental pipeline isolating host inhibitory neurons from transplanted adult mouse visual cortex (V1). Host inhibitory neurons (green) were genetically-labeled via Cre-dependent fluorescent tagging. Embryonic inhibitory neurons (red, MGE or LGE) were transplanted into adult V1 using physiological mapping. V1s were dissected out at the peak of (35 DAT) or after (70 DAT) the transplant-reactivated critical period. Cell populations were separated by FACS and prepared for sequencing. Each sample n= 2-3 V1 replicates pooled together. 35 DAT MGE: n= 8 samples, 19 mice. 35 DAT LGE: n= 5 samples, 10 mice. 35 DAT adult no tx: n=7, 18 mice. 70 DAT MGE: n= 4 samples, 10 mice. 70 DAT adult no tx: n= 4 samples, 12 mice. V1 microglia collected as an additional pipeline control. RNAseq data was collected in 3 batches. Differential expression (DE) analysis conducted via DESeq2. Gene ontology annotation of enriched **(B)** biological processes (GO Biological Process database), **(C)** cellular components (GO Cellular Components database), and **(D)** brain regions (Allen Brain Atlas Coexpression database) for the top 1% (by ranked p-value) of MGE vs. LGE DE genes (∼300). Analysis conducted via ShinyGO 0.77, hits ranked by FDR. Gene ontology annotation of predicted transcription factor interactions for the top 1% (by ranked p-value) of MGE vs. LGE DE genes (∼300) using the **(E)** ENCODE and ChEA Consensus TFs from Chip-X database, and **(F)** TRANSFAC and JASPAR PWMs database. **(G)** Volcano plot of the top 1% (by ranked p-value) of MGE vs. LGE DE genes (∼300). Red points represent significantly upregulated genes, blue points represent significantly downregulated genes. Grey points represent genes with a smaller fold change (<1 or > −1).

Donor inhibitory neurons were isolated from microdissected embryonic (E13.5-14.5) brain regions, either: a) the medial ganglionic eminence (MGE), which produces parvalbumin-(PV) and somatostatin-(SOM) expressing inhibitory neurons and whose transplantation reactivates plasticity, or b) the lateral ganglionic eminence (LGE), which also produces PV and SOM subpopulations but whose transplantation does not alter plasticity (*2–3*). Each transplant cohort included both LGE transplant controls to account for cell transplantation-related gene expression, and non-transplanted, age-matched controls to account for baseline age-related gene expression. (Fig 1B). Transplant cohorts were collected at timepoints that represented two distinct physiological states – either: a) 35 days after transplantation, during the well-documented peak of transplant-reactivated plasticity (35 DAT), or b) 70 days after transplantation, after closure (70 DAT) (Fig 2) (*2-4*). V1s were dissected solely from the right hemisphere, including in non-transplanted mice, to preclude variation from hemispheric asymmetry. Due to the small size of V1 (∼ 1mm^3^), tissue was pooled across mice (n= 2-3 V1s each) prior to dissociation. Samples were then separated by cell population using fluorescence-activated cell sorting and processed for RNA sequencing.

**Figure 2.**
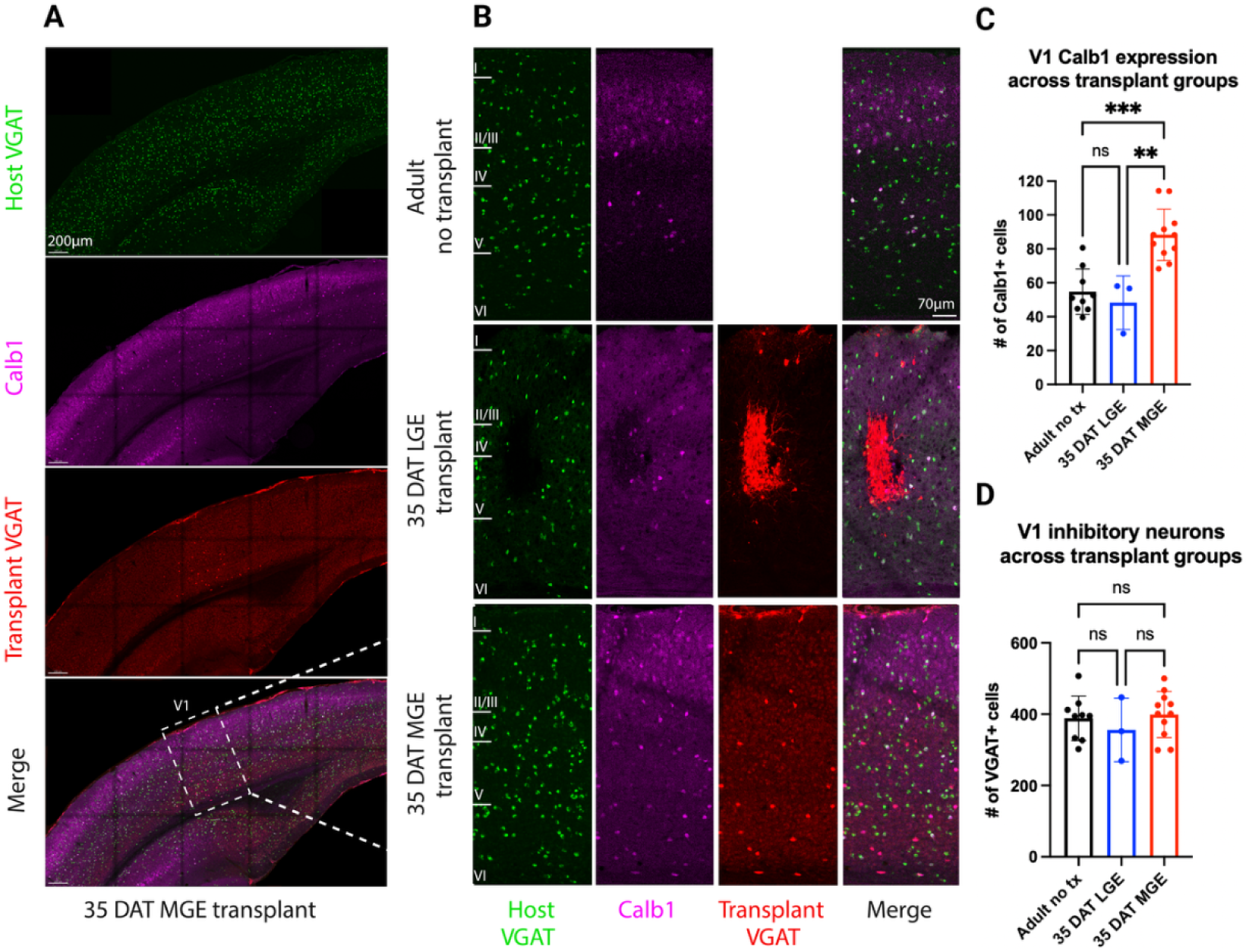
MGE transplantation upregulates Calb1 expression. (A) Example V1 coronal section from adult MGE recipient 35 days after transplantation (DAT). Host inhibitory neurons (Host VGAT) genetically labeled by VGAT-ZsGreen fluorescence. Transplanted inhibitory neurons (Transplant VGAT) genetically labeled by VGAT tdTomato fluorescence. Calb1 protein (purple) labeled by antibody immunostaining. Transplanted inhibitory neurons do not migrate past V1 boundaries. Calb1 upregulation appears restricted to the transplanted region. (B) Magnified V1 sections from adult non-transplanted mice, LGE recipients 35 DAT, and MGE recipients 35 DAT. Cortical layers annotated to the left of host VGAT images. LGE-derived inhibitory neurons remain at the engraftment site. MGE-derived inhibitory neurons appear to populate deeper cortical layers after transplantation. (C) Quantification of Calb1+ cells across transplant groups by animal. Each animal was sampled using 2-4 sections from anterior to posterior V1. V1 bounds for each section was determined using the Allen Brain Atlas. Adult no transplant (no tx) mice (black): n= 9. LGE recipients (blue): n= 3. MGE recipients (red): n= 11. Calb1 levels are upregulated in MGE recipients compared to LGE recipients (MGE vs. LGE p= 0.0011, one-way ANOVA with Turkey’s) and non-transplanted mice (MGE vs. no tx p= 0.0001, one-way ANOVA with Turkey’s). No significant difference between Calb1 levels in LGE recipients and adult non-transplanted mice. Bars are mean +/- SEM. (D) Quantification of VGAT+ cells across transplant groups by animal. Differential Calb1 expression does not appear to be due to significantly different numbers of V1 inhibitory neurons (one-way ANOVA with Turkey’s). Bars are mean +/- SEM.

### Inhibitory neuron transplantation does not activate microglial response

Next, we performed transcriptional analysis in microglia, which are highly reactive to changes in their environment (*10*), and as such, provide a sensitive readout for generic effects of our experimental design on cell physiology. Microglia were fluorescently tagged after V1 dissociation using Cd11b-APC (microglial marker commonly used for flow cytometry) and sorted. Across experimental groups, microglial profiles from MGE, LGE, and non-transplanted adult V1 all showed relatively homogenous expression of homeostatic markers (*11*) (Supp. Fig 1A). Gene ontology (GO) annotation of the most robust DE gene set, microglia from MGE recipients (reactivated plasticity) vs. microglia from LGE recipients (normal), did show upregulation of angiogenesis-related terms (Supp. Fig 1B). Aside from this exception, the signature of homeostatic microglial markers were not substantially altered by inhibitory neuron transplantation or the collection process.

### Host inhibitory neurons upregulate synaptic plasticity components (peak plasticity vs. after closure)

To survey changes resulting from MGE transplantation, we first conducted a broader-scope DE analysis of host inhibitory neurons from MGE recipients 35 DAT vs. host inhibitory neurons from MGE recipients 70 DAT. GO annotation of biological processes showed a strong signature of cell-cell signaling, synaptic transmission, and ion transport, as well as enrichment of extracellular matrix elements, altogether suggestive of neurite outgrowth and synaptic plasticity (Supp. Fig 2A-B). Of note, transcription factor binding analysis confirmed involvement of CREB-related pathways, a known mediator of activity-dependent gene expression with strong ties to plasticity, critical period development, and inhibitory neuron transplantation (*12*) (Supp. Fig 2C).

### Host inhibitory neurons downregulate inhibitory fate markers (reactivated plasticity vs. non-transplanted baseline)

Next, we conducted a finer-resolution DE analysis of host inhibitory neurons from MGE recipients 35 DAT vs. host inhibitory neurons from non-transplanted, age-matched controls 35 DAT. GO enrichment analysis revealed upregulation of immune response components and strong detection of exogenous factors from another organism (Supp. Fig 3A). This finding was also observed in DE analysis of host inhibitory neurons from LGE recipients 35 DAT vs. host inhibitory neurons from non-transplanted, age-matched controls 35 DAT (Supp. Fig 3B). Given the similar GO enrichment, it may suggest that host inhibitory neurons, regardless of the type of transplant, detect the presence of the transplanted cells.

The DE genes in MGE vs. non-transplanted controls were also associated with strong receptor activity and enzyme regulation, further suggesting that host inhibitory neurons experience elevated paracrine signaling (Supp. Fig 3C). Transcription factor enrichment analysis highlighted TFAP2A (regulator of neural development and migration) (*13*), and Runx2 (implicated in inhibitory neuron fate specification) (*14*) (Supp. Fig 3E). Segmentation of DE genes into up- and downregulated gene sets revealed consistent downregulation of genes involved in neuronal fate commitment (Supp. Fig 3E). Network analysis further highlighted specification factors, such as Dlx genes, that hold critical roles in GABAergic differentiation during normal embryonic development (*15*).

### Host inhibitory neurons upregulate Calb1 (reactivated plasticity vs. cell transplant control)

Finally, as a biological screen for changes specific to transplant-reactivated cortical plasticity, we conducted DE analysis of host inhibitory neurons from MGE recipients 35 DAT vs. host inhibitory neurons from LGE recipients 35 DAT. GO annotation of biological processes again showed enrichment of cell-to-cell signaling and ion transport, but also nervous system development mechanisms (Fig 1B). This pattern continued to be observed in the top enriched cellular components with cerebral cortex and developmental brain structures (Fig 1D), and transcription factor enrichment analysis with SUZ12/JARID2 (transcriptional repressor essential for differentiation) (*16*) and EZH2/BMI1 (histone modifier/factor that maintains stemness) (*17*) (Fig 1F).

Among the genes upregulated in host inhibitory neurons from MGE recipients 35 DAT, Calb1 stood out as a potential key molecule for driving cortical plasticity. (Fig 1G). Known for its potent Ca^2+^-binding properties (*18*), Calb1 is a mobile, cytosolic protein that localizes to the soma and dendrites, predominantly within inhibitory neurons (*19*). Loss of Calb1 impairs multiple aspects of synaptic plasticity, such as proper LTP maintenance, paired-pulse facilitation, and neuronal excitability (*20*)– ultimately manifesting as deficits in learning and memory, motor coordination, and sensory integration (*21*). Through its protein interactions, Calb1 also plays non-canonical roles such as suppression of apoptosis (*22*), promotion of neurite outgrowth, and neuronal differentiation (*23*).

Across the central nervous system, Calb1 expression increases during embryogenesis, peaks during postnatal development, and sharply downregulates upon adulthood (*24*). Interestingly, Calb1 is co-expressed by both MGE-derived subpopulations, PV and SOM (*19*), that have been strongly implicated in critical period regulation and inhibitory neuron transplantation (*25*). As such, Calb1 appeared uniquely positioned at the intersection of our DE results and GO analysis.

### Calb1 expression is high during juvenile critical period and transplant-reactivated plasticity

For independent, protein-level validation, we first examined Calb1 expression in adult V1 sections from MGE, LGE, and non-transplanted mice 35 DAT. In MGE recipients, Calb1 expression was significantly higher than in both LGE recipients and non-transplanted adults (Fig 2B). LGE transplantation did not significantly alter Calb1 levels from the non-transplanted baseline (Fig 2C). Similar inhibitory neuron counts across groups (transplant and host) indicates that the increase in Calb1 expression does not stem simply from the addition of Calb1+ transplanted cells (Fig 2D). Changes in Calb1 expression have been previously noted following neuronal transplantation into other brain circuits including in the engrafted cells themselves (*26–27*). Consistent with previous studies, non-transplanted adults showed distinct laminar enrichment in cortical layers II/III and V with segregation into strongly- and weakly-expressing Calb1+ cells (*28*). In contrast, MGE transplantation appears to increase and redistribute Calb1 expression in deeper layers (Fig 2B).

Given previous findings of a shared mechanism between juvenile and transplant-reactivated plasticity (*7*), we examined Calb1 expression in V1 sections from non-transplanted juvenile mice. We assessed three different key timepoints for visual development: P16 (two days after eye-opening and the onset of visual experience), P28 (peak of the juvenile critical period), and P35 (closure of the juvenile critical period) (*29*) (Fig 3A). We found that Calb1 expression peaks at P28 and begins to downregulate in deeper layers at P35 (Fig 3B). Notably, Calb1 levels after MGE transplantation appeared similar to P28 levels (Fig 3C), suggesting that MGE transplantation restores low adult Calb1 expression to higher juvenile levels.

**Fig 3.**
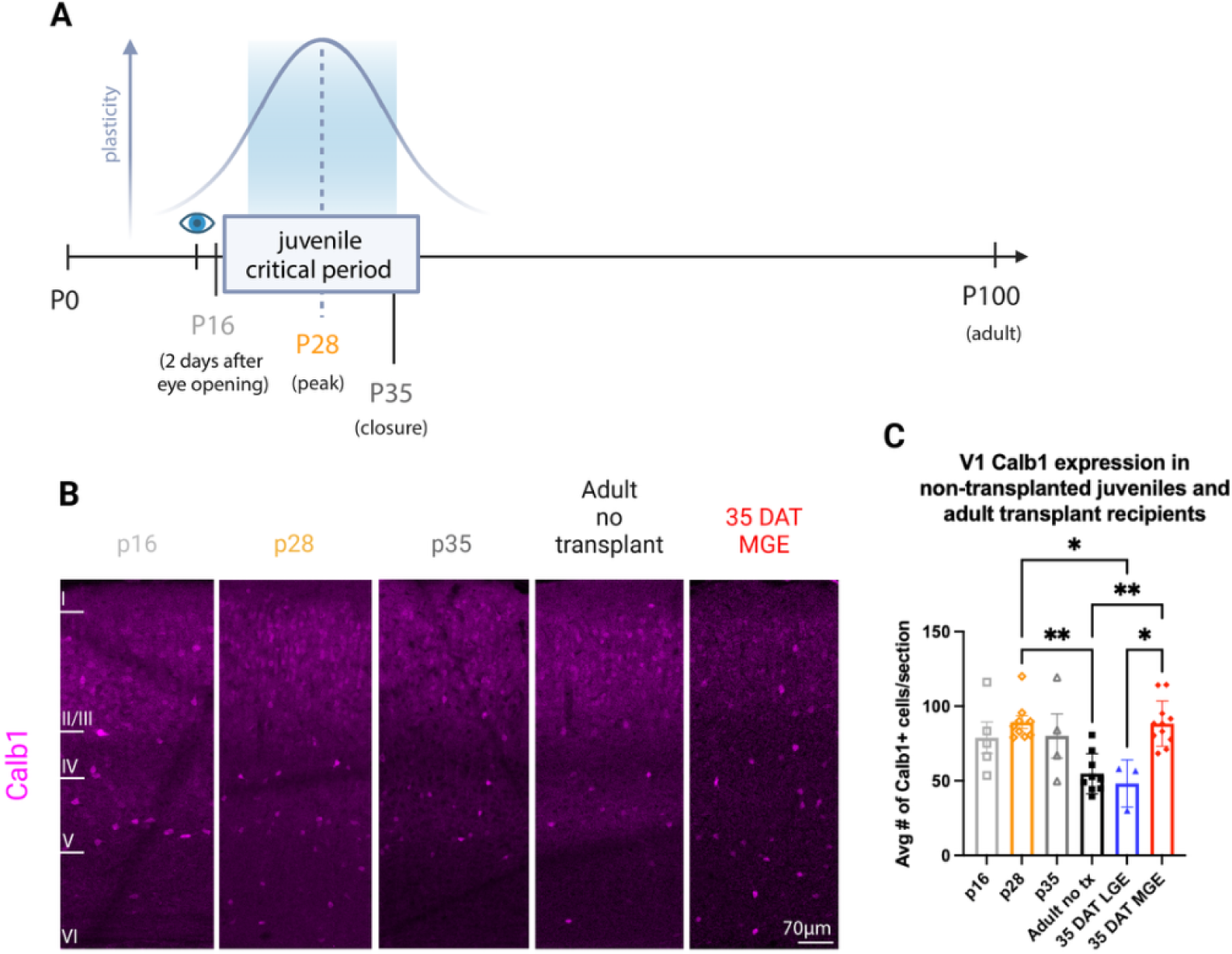
Calb1 expression peaks during the visual critical period. (A) Schematic of assessed timepoints during postnatal visual development. P14, eye opening and onset of visual experience. P16, 2 days after eye opening. P28, peak of the juvenile critical period. P35, beginning of critical period closure. p100, adulthood. Representative curve indicating increasing plasticity with onset of the critical period, and decreasing plasticity upon closure. **(A)** Magnified visual cortex coronal sections from non-transplanted p16 (2 days after eye opening), p28, p35, and adult mice. Example V1 section from MGE recipient 35 DAT included for comparison. Calb1 protein (purple) labeled by antibody immunostaining. CCortical layers annotated to the left of p16 image. **(C)** Quantification of Calb1+ cells across groups by animal. Each animal was sampled using 3-6 sections from anterior to posterior V1. V1 bounds and cortical layers for each section were determined using the Allen Brain Atlas. p16 (light gray): n= 5. p28 (orange): n= 9. p35 (dark gray): n=4. Adult no tx (black): n= 9. LGE recipients (blue): n= 3. MGE recipients (red): n= 11. Calb1 levels increase during the peak of the juvenile critical period, and decline with age (p28 vs. adult no tx: p= 0.0019, one-way ANOVA with Turkey’s). Adult Calb1 levels re-upregulate upon MGE transplantation (Adult no tx vs. MGE: p= 0.0015, one-way ANOVA with Turkey’s). MGE vs. LGE: p=0.0124, p28 vs. LGE: p=0.0123 (one-way ANOVA with Turkey’s). Bars are mean +/- SEM. Non-significant stats omitted for visualization.

### Ocular dominance plasticity is diminished after Calb1 protein knockdown in V1 inhibitory neurons

As a preliminary test of whether Calb1 could be functionally linked to critical period plasticity, we knocked down Calb1 protein expression in V1 inhibitory neurons and assessed critical period cortical plasticity. The null hypothesis was that plasticity would remain unchanged after Calb1 knockdown. To selectively knock down Calb1 in V1 interneurons, we physiologically mapped binocular V1 in VGAT-Cre mice and injected a Cre-dependent, Calb1 shRNA-expressing virus (AAV1-CAG-DIO-mCalb1-shRNAmir) (Fig X). To probe normal critical period plasticity, VGAT-cre pups (p3-5) were unilaterally-injected with either the shRNA AAV or saline. Knockdown of Calb1 protein in cells infected with GFP reporter was indicated by weaker anti-Calb1 immunofluorescence compared to surrounding uninfected interneurons, other-hemisphere V1, and saline-injected controls (Fig 4C).

**Fig 4.**
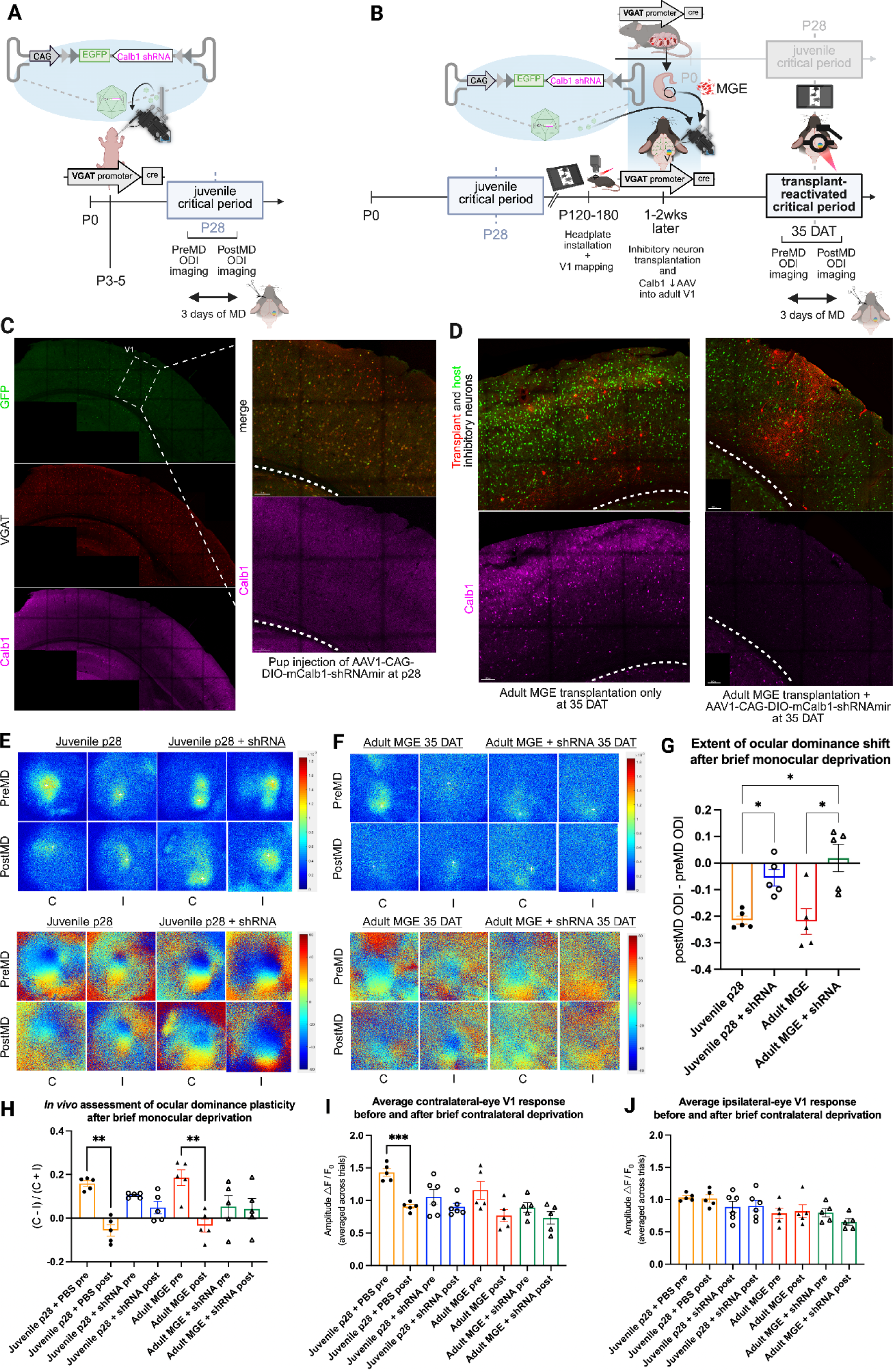
Reduction of Calb1 activity in V1 inhibitory neurons diminishes expression of ocular dominance plasticity. (A) Schematic of experimental design for assessing effects of Calb1 inactivation on juvenile experience-dependent plasticity. Pup V1 is targeted stereotactically. **(B)** Schematic of experimental design for assessing effects of Calb1 inactivation on transplant-reactivated plasticity. Adult V1 is physiologically targeted via intrinsic signal optical imaging. AAV construct restricts Calb1 inactivation to Cre-expressing inhibitory neurons. **(C)** Example V1 coronal section from non-transplanted juvenile (p28) mice expressing Calb1 shRNA in inhibitory neurons. Viral GFP reporter in green, Calb1 antibody labeling in magenta, genetically-labeled inhibitory neurons in red. Virus expression and reduction in Calb1 expression/intensity remains focal to the injection site. Magnified views show successful infection of inhibitory neurons (yellow cells) and strong reduction of Calb1 expression. Reduction in Calb1 expression does not spread from injected V1 to adjacent cortical regions or hippocampus below. **(D)** Example V1 coronal section from adult mice receiving MGE transplantation only (left panels) vs example V1 coronal section from adult mice receiving MGE transplantation and Calb1-silencing AAV (right panels). MGE transplantation strongly upregulates Calb1 expression. Reduction in Calb1 expression does not spread to hippocampus below. Example contralateral (deprived eye, C) and ipsilateral (open eye, I) response amplitudes before and after 3 days of monocular deprivation (MD) in **(E)** juvenile and **(F)** adult-transplanted injection groups. **(G)** Ocular dominance shifts across injection groups. Shifts are calculated: postMD ODI-preMD ODI. shRNA-AAV recipients at peak plasticity timepoints show reduced shifts in ocular dominance after brief monocular deprivation compared to saline-injected mice (p28 + PBS vs. p28 + Calb1 shRNA: p= 0.0174; MGE vs. MGE + Calb1 shRNA: p= 0.0494, Brown-Forsythe and Welch ANOVA with Dunnett’s). Bars are mean +/- SEM. **(H)** Ocular dominance indices (ODI) before and after MD. ODIs are calculated: (C-I)/(C+I). Juvenile (p28) mice and MGE-transplanted adult mice exhibit significantly different pre and postMD ODIs after brief monocular deprivation, as previously reported (p28 + PBS preMD vs. p28 + PBS post MD: p= 0.0011; adult MGE preMD vs. adult MGE postMD: p= 0.0056, Brown-Forsythe and Welch ANOVA with Dunnett’s). Pre and postMD ODIs for PBS-injected mice are not significantly different (p28 + shRNA preMD vs. p28 + shRNA postMD: p= 0.3538, adult MGE + shRNA preMD vs. adult MGE +shRNA postMD: p= 0.9997, Brown-Forsythe and Welch ANOVA with Dunnett’s). Bars are mean +/- SEM. **(I-J)** Quantification of contralateral and ipsilateral response amplitudes before and after brief monocular deprivation. Response amplitudes are an average of 3-4 recordings for each eye, alternating between eyes. After 3 days of MD, PBS-injected p28 mice show a significant reduction in contralateral responses from pre to postMD (p= 0.00008, Brown-Forsythe and Welch ANOVA with Dunnett’s). A similar contralateral reduction, though not statistically significant in this sampling, is found in MGE-transplanted adult mice (p=0.172, Brown-Forsythe and Welch ANOVA with Dunnett’s). Non-significant comparisons omitted for visualization.

We assessed cortical plasticity in V1 using eye-specific cortical mapping of ocular dominance. Mice are normally contralateral-eye dominant, but disruptions in visual experience, such as monocular deprivation (MD), can shift ocular dominance to the remaining ipsilateral-eye (*1*). Importantly, ocular dominance plasticity (ODP) is predominantly restricted to juvenile mice, as normal adult mice express a distinctively weaker and shorter-lasting capacity for circuit reorganization (*31*). Two days before the peak of the normal critical period (∼P26), juvenile injected mice received headplates and baseline V1 responses were recorded from each eye using ISOI. Contralateral and ipsilateral responses were then used to calculate an ocular dominance index (ODI), a measure of eye-specific response strength relative to each other. Immediately following baseline imaging, the eye contralateral to the injected hemisphere was sutured shut and deprived of visual stimuli through the peak of the ocular dominance critical period (p28). After 3 days of MD, eyelid sutures were removed and V1 responses recorded from each eye again. Calb1 knockdown in juveniles reduced baseline contralateral response, which did not further reduce upon brief MD. As compared to saline-controls, shRNA knockdown of Calb1 in V1 inhibitory neurons resulted in negligible ocular dominance plasticity, consistent with the importance of Calb1 in cortical plasticity.

We similarly assessed transplant-reactivated plasticity after Calb1 knockdown. Adult VGAT-cre mice received either co-injection of donor MGE with shRNA AAV or donor MGE only. Calb1 shRNA resulted in knockdown of Calb1 immunofluorescence intensity compared to MGE-only controls (Fig 4D). A few days before the peak of the reactivated critical period (∼33 DAT), eye-specific V1 responses were recorded using ISOI and baseline ODIs calculated. Adult mice then underwent brief MD (3 days) and eye-specific V1 responses measured again. Consistent with previous studies, MGE transplanted adults demonstrate a significant shift in ocular dominance after brief MD, facilitated by specific reduction of the contralateral eye response strength (Fig 4I). In contrast, MGE/shRNA-injected adults had ODIs that were not significantly different after MD (Fig 4H), suggesting impairment of transplant-reactivated ocular dominance plasticity after Calb1 knockdown.

The results that ocular dominance plasticity is diminished after Calb1 knockdown were intriguing and consistent with the importance of Calb1 in the regulation of cortical plasticity. However, the AAV injections could have interfered with plasticity by mechanisms other than shRNA-induced protein knockdown e.g., by AAV or shRNA toxicity. We note that transplanted inhibitory neurons do not disperse as usual when co-injected with the shRNA AAV (Fig 4D). While these results do not provide definitive proof of necessity, they invite exploration of Calb1’s role within inhibitory neurons for the regulation of cortical plasticity.

### Upregulation of Calb1 in adult V1 inhibitory neurons reactivates juvenile plasticity

Given the observed correlations between Calb1 expression and cortical plasticity, we tested whether directly upregulating Calb1 in adult V1 reactivates juvenile plasticity. Previous *in vivo* studies showed that reinstatement of Calb1 expression in hippocampal circuits could rescue cognitive deficits imposed by its suppression (*30*). To upregulate Calb1 specifically within V1 inhibitory neurons, we designed a Cre-dependent Calb1-overexpressing virus (AAV1-CMV-DIO-mCalb1-T2A-eGFP) and injected it into mapped binocular V1 (Fig 5A).

**Fig 5.**
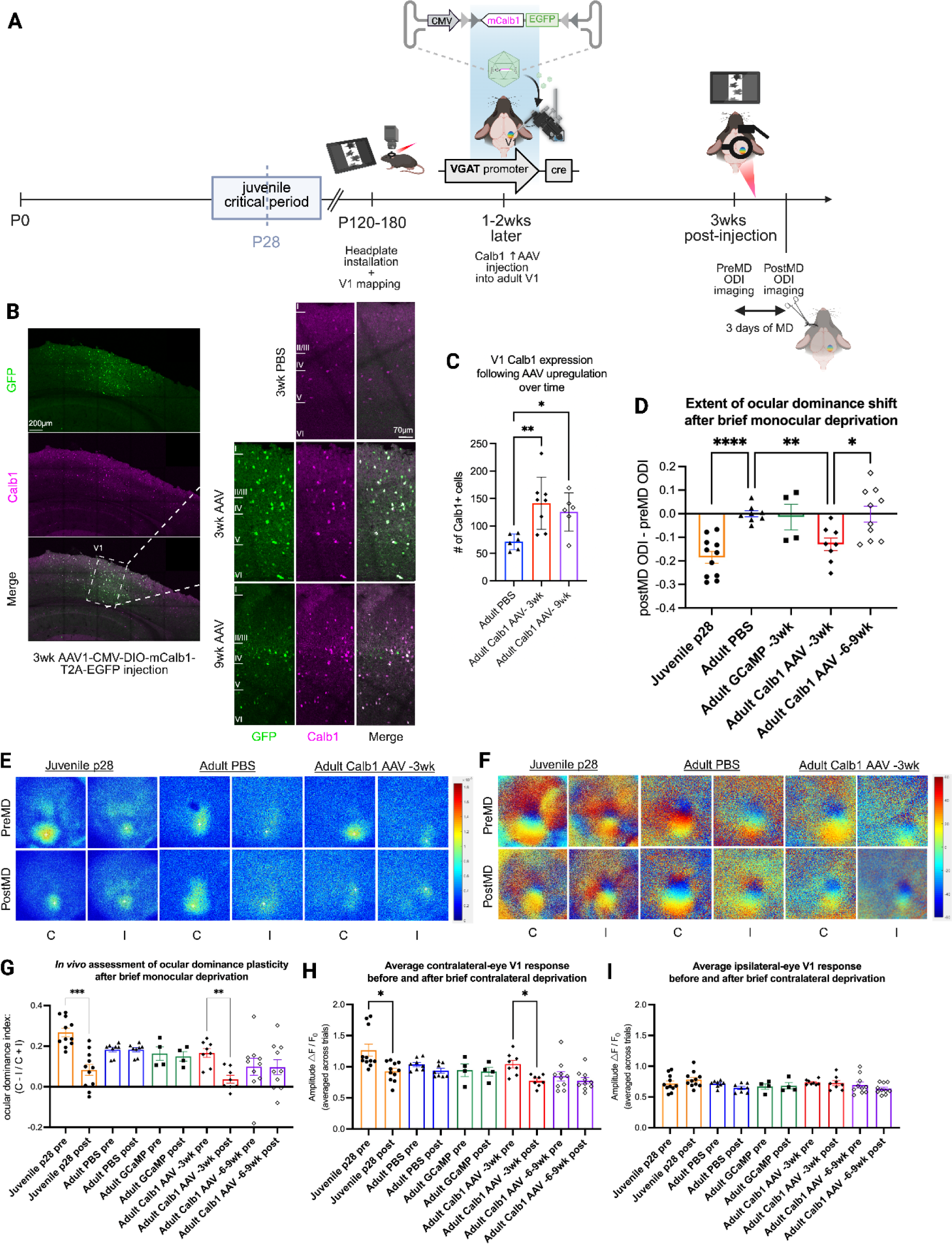
Calb1 re-upregulation in adult V1 inhibitory neurons restores juvenile ocular dominance plasticity. (A) Schematic of experimental design. AAV construct restricts Calb1 overexpression to inhibitory neurons. **(B)** Example V1 coronal section from adult mouse 3 weeks after AAV injection. Successful integration labeled by GFP (green). Calb1 (purple) upregulation appears restricted to the injected region. Magnified V1 sections from adult saline-injected mice (3wk PBS) and AAV recipients 3 weeks post-injection (3wk AAV). Cortical layers annotated on the leftmost images. **(C)** Quantification of Calb1+ cells across injection groups by animal. Each animal was sampled using 2-4 sections from anterior to posterior V1. V1 bounds for each section was determined using the Allen Brain Atlas. 3wk PBS (black): n= 6. 3wk AAV (red): n=8, 9wk AAV (purple): n=6. Calb1 levels are upregulated in AAV-injected adult mice compared to PBS-injected adult mice at 3 weeks and 9 weeks post (PBS vs. AAV −3wk: p= 0.01; PBS vs. AAV −9wk: p= 0.026, Brown-Forsythe and Welch ANOVA with Dunnett’s). Bars are mean +/- SEM. **(D)** Ocular dominance shifts across injection groups. Shifts are calculated: postMD ODI-preMD ODI. Juvenile p28 mice demonstrate significant shifts in ocular dominance after brief MD compared to adult mice (p28 vs. adult PBS: p= 0.0002, Brown-Forsythe and Welch ANOVA with Dunnett’s). While inhibitory neuron-specific expression of GCaMP does not alter plasticity from adult baseline (adult PBS vs. adult GCaMP: p>0.9999), AAV recipients at 3 weeks post-injection show significant shifts in ocular dominance after brief MD in comparison (adult AAV vs. adult PBS: p= 0.017, Brown-Forsythe and Welch ANOVA with Dunnett’s). OD shifts measured beyond 6 weeks post-injection diminished compared to measurements at 3 weeks (AAV-3wk vs. AAV-6-9wks: p= 0.0441, Brown-Forsythe and Welch ANOVA with Dunnett’s). Non-significant comparisons omitted for visualization. Bars are mean +/- SEM. **(E-F)** Example contralateral (deprived eye, C) and ipsilateral (open eye, I) response amplitudes before and after 3 days of monocular deprivation (MD) for each adult injection group, as well as unaltered juvenile positive controls (p28). **(H)** Ocular dominance indices (ODI) before and after MD. ODIs are calculated: (C-I)/(C+I). After 3 weeks, AAV-injected mice exhibit significantly different pre and postMD ODIs (AAV-3wk preMD vs. AAV-3wk post MD: p= 0.0026, Brown-Forsythe and Welch ANOVA with Dunnett’s). Pre and postMD ODIs for PBS-injected adult mice are not significantly different (adult PBS preMD vs. adult PBS postMD: p>0.9999, Brown-Forsythe and Welch ANOVA with Dunnett’s) while pre and postMD ODIs for p28 juveniles are significantly different as expected (p28 preMD vs. p28 postMD: p= 0.0002, Brown-Forsythe and Welch ANOVA with Dunnett’s). Adult baselines are slightly more binocular than juvenile baselines (p28 preMD vs. adult PBS preMD: p= 0.0163, Brown-Forsythe and Welch ANOVA with Dunnett’s). Non-significant comparisons omitted for visualization. Bars are mean +/- SEM. **(H-I)** Quantification of contralateral and ipsilateral response amplitudes before and after brief monocular deprivation. Response amplitudes are an average of 3-4 recordings for each eye, alternating between eyes. After 3 weeks, AAV-injected mice show a significant reduction in contralateral responses from pre to postMD (adult AAV preMD vs. adult AAV postMD: p= 0.0147, Brown-Forsythe and Welch ANOVA with Dunnett’s), similar to unaltered p28 juvenile mice (p28 preMD vs. p28 postMD: p=0.0393, Brown-Forsythe and Welch ANOVA with Dunnett’s). Non-significant comparisons omitted for visualization.

Successful upregulation of Calb1 protein within targeted cells and localization of the GFP reporter to V1 was determined at 2 weeks post-injection during initial virus optimization and confirmed after each experiment via immunohistochemistry (Fig 5B-C). Upregulation of Calb1 protein remained specific to the virus injection site, where adjacent cortical regions continue to exhibit the reduced adult Calb1 expression pattern (Fig 5B). The majority of AAV-infected inhibitory neurons were co-labeled with anti-Calb1, although some infected neurons show weaker or no immunofluorescence. Similar to P28 and MGE 35 DAT, AAV-mediated upregulation of Calb1 approximately doubles the number of Calb1-expressing cells and redistributes higher-expressing cells across deeper layers (Fig 5C).

ODP was assessed in juvenile mice or adults unilaterally-injected with Calb1 AAV, saline, or a control AAV driving expression of a different calcium buffer and fluorophore, GCaMP6s. (AAV1-CMV-DIO-GCaMP6s). Baseline imaging began 3 weeks after injections, prior to the expected period of reactivated cortical plasticity (2). Immediately following baseline imaging, the eye contralateral to the injected hemisphere was sutured shut and deprived of visual stimuli for 3 days, similar to the Calb1 knockdown experiments above.

Similar to saline-injected adult mice, GCaMP-injected mice did not exhibit an ODP shift (Fig 5D), indicating that cre-dependent upregulation of the calcium buffer calmodulin fused to GFP in adult V1 inhibitory neurons does not induce cortical plasticity. Remarkably, Calb1 AAV-injected adult mice demonstrate a significant reduction of contralateral eye visual input to V1, a hallmark of juvenile ocular dominance plasticity in mice (Fig 5H; 31). This pattern is expected in juvenile mice, while ocular dominance shifts in adult mice typically involve a gain in response from the non-deprived eye (31). Similar to the normal juvenile critical period (P16-P35, Fig 3B), and transplant induced critical period reactivation (25DAT-43 DAT, 2), Calb1-reactivated plasticity was found to be reduced 3 weeks later and on (Fig 5D), suggesting that endogenous brake mechanisms may similarly engage upon AAV-mediated Calb1 upregulation.

## DISCUSSION

Our findings reveal that transplanted embryonic inhibitory neurons reactivate a developmental program within adult host inhibitory neurons. The upregulation of membrane-bound proteins capable of nuclear translocation, chromatin remodeling elements, and positive transcriptional regulators offer converging evidence that host inhibitory neurons undergo a cell-state transition. The upregulation of Calb1, a high-capacity Ca^2+^-binding protein with dual buffer and sensor capability (*18*), may enable adult inhibitory neurons to once again rapidly respond to changes in sensory input. Apart from potently altering signal transduction, Calb1 upregulation has also been closely linked to neurotrophin-mediated differentiation and maturation of post-mitotic neuronal progenitors (*32–33*).

Given the modest number of transplanted inhibitory neurons needed to reactivate cortical plasticity, how do signals from the transplanted cells rejuvenate an entire circuit? While some host inhibitory neurons may receive factors directly from the transplanted cells, it may be that others receive those signals from their newly rejuvenated neighbors. Moreover, if transplanted inhibitory neurons secrete factors while they migrate through the host tissue, it may be that those factors can affect other host brain cell-types. For example, while upregulation of angiogenesis in adult microglia during transplant-reactivated plasticity may be a response to the increased metabolic demand brought on by neuronal rejuvenation, it may also represent the return of microglia to a juvenile state (*34–35*).

Despite the reactivation of embryonic patterns of gene expression, transplant-rejuvenated inhibitory neurons do not appear to functionally dedifferentiate. Their retention of neuronal morphology (*2,36*), synapse-related gene expression, and visual properties (*3,7,45*) suggests instead a more restricted form of cellular plasticity where adult neurons are restored to an earlier developmental state without more extensive reprogramming. This effect is reminiscent of heterochronic parabiosis, where older endogenous cells are signaled to “age-match” younger exogenous cells (*37*). Interestingly, Calb1 expression is significantly restored by heterochronic parabiosis in an Alzheimer’s mouse model (*38*). Taken together, Calb1 may be a robust biomarker of neuronal rejuvenation.

Studies of juvenile critical periods have focused primarily on the role of PV inhibitory neurons (*1*). However, PV inhibitory neurons are highly heterogenous (*39*) and encompass nearly 40% of all inhibitory neurons. Interestingly, transplantation of MGE-derived SOM inhibitory neurons also reactivates plasticity (*40*)-suggesting that the factors responsible for transplant-reactivated plasticity are not exclusive to PV inhibitory neurons, but in fact shared across PV and SOM subpopulations. Given that Calb1 is transiently increased in both PV and SOM subpopulations during early development (*19*), its upregulation may represent a property of inhibitory neurons that more precisely marks critical periods.

The clinical application of inhibitory neuron transplantation is limited by numerous technical challenges including inefficient cell manufacturing, restricted biodistribution, and safety risks such as immune rejection. As an alternative to inhibitory neuron transplantation (*41*), here we show initial proof-of-principle that transplant-reactivated cortical plasticity can be recapitulated without a cellular vehicle. It may be that, much like other cellular reprogramming methods (*42*), optimal reactivation requires a set of rejuvenation factors yet to be discovered. Targeted molecular rejuvenation offers a new avenue for therapeutic repair for a broad spectrum of neurological disorders stemming from synaptic dysfunction.

## Acknowledgments

We thank the UCI Genomic Research and Technology Hub, UCI Center for Neural Circuit Mapping, UCI Flow Cytometry Core, UCI Optical Biology Core, and J. Zeitoun for ISOI programming assistance.

## Supplementary Materials

## Materials and Methods

### Surgical Procedures

All protocols and procedures followed the guidelines of the Animal Care and Use Committee at the University of California, Irvine. All animals (C57BL/6 background) are raised on 12-hour light/dark cycles, with food and water available ad libitum. Breeders are housed with no more than two other littermates per cage. Although previous studies do not report significant sex differences in transplant-reactivated plasticity, all cohorts of experimental mice included near equal numbers of males and females.

To label host inhibitory neurons, adult mice with green fluorescent inhibitory neurons were generated by crossing homozygous VGAT-cre females (JAX 028862) with cre-dependent ZsGreen males (Ai6, JAX 007906). Resulting litters were aged to P120-P180 (4-6mo) prior to experimental onset. To distinguish transplanted inhibitory neurons from host inhibitory neurons, donor embryos with red fluorescent inhibitory neurons were generated by crossing homozygous VGAT-cre females with cre-dependent tdTomato males (Ai14, JAX 007914). Viral injections were conducted in hemizygous VGAT-tdTomato recipient mice. Colony genotypes were periodically confirmed by PCR (Transnetyx), and successful cre recombination was individually confirmed before each experiment via fluorescent reporters. To ensure stability for downstream intracerebral procedures, custom-printed headplates were affixed to the skulls of adult recipient mice. Mice received a subcutaneous injection of analgesic Carprofen (0.8 mg/ml; Rimadyl), supplemental hydration (Lactated Ringer’s), and eye ointment (Systane) prior to surgery. Body temperature was maintained throughout the procedure using a feedback-controlled heating pad. After anesthetizing with 2% isoflurane, ear bars were fitted to temporarily stabilize head axis. First, local topical and injectable lidocaine were administered, then intact skull was surgically exposed and reinforced with Vetbond (3 M, Vetbond, 1469SB). V1-adjacent regions were additionally reinforced with a layer of dental acrylic (Lang Ortho-Jet Powder and Ortho-Jet Powder Liquid). Custom-printed headplates were then affixed using dental acrylic, with the 4-5mm window centered over V1. Lastly, V1 windows were sealed with a final layer of Vetbond, and mice returned to heated home cages upon waking.

For consistency, all headplates were installed by a single experimenter, on the right hemisphere only. Mice received subcutaneous injections of Carprofen to reduce pain and inflammation (up to 3 days) after installation. Previous studies suggest that social enrichment may enable ocular dominance plasticity in adult mice (*43*). To avoid confounding factors, all experimental animals were single-housed after headplate installation.

### Intrinsic Signal Optical Imaging (ISOI) of Visual Cortical Responses

To precisely target V1, the central, binocular region (bV1) was physiologically determined for each individual mouse using ISOI. After inducing and maintaining a steady plane of light anesthesia with 0.8-1.2% isofluorane, headplated mice were secured into the imaging rig. Headplate windows were filled with PBS and covered with a 10mm glass coverslip. PBS was added as needed throughout sessions to keep coverslips level with the top of the headplate. Eye moisture was maintained using periodic silicone oil application, and body temperature regulated by a feedback-controlled heating pad.

Intrinsic signal images were collected using a custom-designed macroscope (Nikon 135 × 50 mm lenses) equipped with a camera (Dalsa 1M30 CCD) positioned over the headplate. First, a green (530 nm) light-emitting diode (LED) was used to visualize and capture an image of surface vasculature. Then, the camera was refocused ∼450-550 μm below the pia surface to target cortical layer II/III of bV1, and a red (617 nm) LED light used to acquire the intrinsic signal. Visual stimuli was displayed on a monitor (Acer V193, 53 × 33cm, 60 Hz refresh rate, 20 cd/m2 mean luminance) positioned 25cm in front of the mice, aligned to their midline and eye line. Stimuli was generated using MATLAB Psychophysics Toolbox extensions and consisted of contrast-modulating sweeping noise restricted to −5° to +15° visual field azimuth and −18° to +36° visual field elevation. Each imaging session was a 5min presentation of stimuli to both eyes. Custom-written Matlab scripts using Fourier analysis were used to generate phase maps of bV1 responses. Resulting phase maps were then overlaid on top of vasculature images to visualize bV1 bounds and determine optimal injection sites.

### Inhibitory Neuron Transplantation for Transcriptomics

To generate donor embryonic tissue, VGAT-cre females were briefly cohoused with cre-dependent tdTomato males overnight (<20hrs). Upon successful breeding, pregnant dams were sacrificed ∼2wks later by isofluorane and cervical dislocation. E13.5-14.5 VGAT-tdTomato embryos were quickly dissected out into chilled L-15 solution (Gibco, 21083027), and forebrains microdissected into a second petri dish of chilled L-15. tdTomato signal was confirmed in each forebrain prior to MGE and LGE microdissection using an epifluorescence scope. Isolated MGE and LGE tissue were then collected in separate tubes of L-15 with 2% (v/v) of DNase I (Roche, 0471672800) and placed on ice.

Headplated adult VGAT-ZsGreen mice were anesthetized with 2% isoflurane and secured into the surgical rig. All recipients received pre-operative Carprofen, Ringer’s, and eye ointment. Body temperature was maintained throughout the procedure using a feedback-controlled heating pad, and anesthesia gradually reduced as needed. To allow the injection micropipette to penetrate, skull underlying mapped bV1 injection sites was thinned using a dental drill (Midwest, 78044) and carbide burr (FG1/4).

Glass micropipettes (Wiretrol 5μl, Drummond Scientific Company) were pulled with ∼75um diameter tips and beveled. To dissociate donor tissue into injectable cell suspensions, isolated MGE or LGE were front-loaded into prepared micropipettes immediately prior to transplantation. Loaded micropipettes were then angled perpendicularly, relative to the brain surface curvature of each individual recipient. Injections into mapped bV1 sites were performed using a custom-made hydraulic manipulator (Narishige, MO-10) between ∼400-600um below the surface at 10nl/min. Each recipient received 100nl per injection for a total of 2 injections (∼2 isolated MGEs/LGEs). After injection, micropipettes were incrementally withdrawn to prevent backflow and sites resealed with Vetbond. Post-operative mice were returned to heated home cages upon waking, and received subcutaneous injections of Carprofen to reduce pain and inflammation (up to 3 days) after installation.

To maintain consistency, experimental agents (MGE, LGE, saline) were injected only into V1 on the right hemisphere. All tissue dissections were conducted by a single experimenter, and all intracerebral transplantations by another single experimenter. Each transplant cohort included balanced numbers of MGE, LGE, and non-transplanted recipients.

### V1 Isolation, Dissociation, Cell Sorting, and RNA Extraction

At their designated experimental timepoints, mice (35 DAT, 70 DAT, non-transplanted littermates) were sacrificed via 1.2% tribromoethanol (Avertin, 18ml/g). After injection, mice were transcardially perfused with RNAase inhibitor (Superase) and transcriptional inhibitors (Actinomycin D, Triptolide) diluted in chilled L-15 solution for 5min. To mitigate cell cycle discrepancies, perfusions for each collection cohort began at the same time of day and were completed within 2hrs. Brains were then dissected out, and V1s isolated using stereotaxic coordinates centered on mapped bV1 (3-4mm lateral, −0.5-0.5mm anterior to posterior bregma). For consistency, all perfusions were conducted by a single experimenter, and V1 dissections conducted by another single experimenter. Isolated V1 tissue was then homogenized into smaller pieces and suspended inadditional L-15/inhibitors solution in gentleMACS C tubes (Miltenyi Biotec). Tissue chunks were further dissociated using the gentleMACS Octo Dissociator with Heaters (Miltenyi Biotec) and Adult Brain Dissociation Kit (mouse and rat, Miltenyi Biotec). Resulting cell suspensions were incubated with microglial antibody (Cd11b-APC, 1:300) and cell viability indicator DAPI for 15min.

Immunolabeled samples were then further refined via fluorescence using UCI’s Institute for Immunology Flow Cytometry Facility (BD FACSAria Fusion Sorter). Following standard FACS screening for cell viability/doublets and fluorescence-gating based on control samples, samples were separated by cell population (ZsGreen-host inhibitory neurons, APC-microglia). All sorted populations were collected into 1ml RNA stabilizer (Trizol) on ice. Finally, RNA was extracted from each sample using the RNA Clean & Concentrator kit (Zymo) and promptly stored at −80C. RNA quality was assessed with a Bioanalyzer (Agilent); only samples with high RNA integrity (RIN >8) were selected for library generation. RNA was quantified using a Qubit (Thermo Fisher Scientific).

### Library Preparation and Sequencing

Stored RNA samples were sequenced in multiple batches. To reduce batch effects, samples across collection cohorts were randomly assigned into batches using a custom script. Libraries were prepared using the SMART-Seq® v4 PLUS Kit (Takara Bio). cDNA quality assessment and quantification was performed with a Bioanalyzer and Qubit, respectively. Only high-quality libraries were sequenced using a Novaseq 6000 (Illumina) for 100bp reads and read depth of 30M per sample.

### Bioinformatic Analyses

FASTQ files were first preprocessed and screened for read quality using FASTQC. The unstranded reads were then pseudoaligned to the mouse GRCm38 index and transcripts quantifed via Salmon (*44*). Low-count genes (reads <10) across samples were filtered out prior to DE analysis using DESeq2. A multi-factor design was applied to account for single vs paired-end sequencing, and batch effects across experimental groups were removed (limma). Despite observed variation within experimental groups, dispersion plots across samples and cell-types were typical, and gene count outliers were not detected.

GO enrichment analyses (i.e. protein-protein interaction, transcription factor binding site) were conducted via ShinyGO using various publicly-available, experimentally-validated/supported databases. Predicted but poorly-annotated (∼270 “Gm-“ and “-Rik”), lowly-expressed (base mean < 5), and smaller effect (log2 fold change between −1 and 1) DE genes were excluded from GO analyses. Resulting GO annotations were filtered to consolidate gene sets with >90% redundancy.

### Immunohistochemistry

Mice were sacrificed via 1.2% tribromoethanol. After injection, mice were transcardially perfused with chilled 4% PFA in PBS for 5min at 6ml/min. Collected brains were post-fixed overnight and cryoprotected with 30% sucrose. Brains were sectioned coronally into 30um free-floating sections using a freezing sliding microtome (Microm, HM450). To reduce batch effects, samples across collection cohorts were randomly assigned into batches using a custom script. To representatively sample V1, sections were selected every ∼200um from anterior to posterior V1. V1 sections were then permeabilized with 0.3% Triton-X in PBS for 1hr, and blocked with 1% normal donkey serum in 0.3% Triton-X/PBS. Following overnight incubation with primary antibodies at 4C, sections were washed 3x in 0.3% Triton-X/PBS and incubated with secondary antibodies for 2hr at room temperature. Sections were again washed 3x in 0.3% Triton-X/PBS, then mounted and coverslipped using Fluoroshield with DAPI (Millipore-Sigma, F6057).

To visualize Calb1 protein expression, sections were incubated with donkey anti-rabbit Calb1 primary antibody (abcam, 1:500), followed by donkey anti-rabbit Alexa 647 secondary antibody (1:500). To visualize transplanted inhibitory neurons, sections were incubated with donkey anti-RFP (Fisher MA5-15257, 1:250) to amplify the VGAT-tdTomato signal, followed by donkey anti-mouse Alexa 568 secondary antibody (1:250). Given the intensity of VGAT-ZsGreen signal, host inhibitory neurons were visualized without additional amplification.

### Intracerebral Viral Injection

To selectively knock down Calb1 in V1 interneurons, we physiologically mapped binocular V1 in VGAT-Cre mice and injected a Cre-dependent, Calb1 shRNA-expressing virus (AAV1-CAG-DIO-mCalb1-shRNAmir). To upregulate Calb1 expression specifically within adult inhibitory neurons, adult non-transplanted VGAT-tdTomato mice were injected with cre-restricted, Calb1-overexpressing virus (AAV1-CMV-DIO-mCalb1-T2A-eGFP, UCI CNCM). An eGFP fluorescent tag was again included in the polycistronic construct to visualize successful AAV incorporation and expression. As a control, we used an AAV driving expression of a different calcium buffer and fluorophore, GCaMP6s. (AAV1-CMV-DIO-GCaMP6s). After headplate installation and V1 mapping, adult VGAT-tdTomato mice were anesthetized with 2% isoflurane and secured into the surgical rig. All recipients received pre-operative Carprofen, Ringer’s, and eye ointment. Body temperature was maintained throughout the procedure using a feedback-controlled heating pad, and anesthesia gradually reduced as needed. To allow the injection micropipette to penetrate, skull underlying mapped bV1 injection sites was thinned using a dental drill and FG1/4 carbide burr.

Glass micropipettes were pulled with ∼75um diameter tips and beveled. Virus was front-loaded into prepared micropipettes and angled perpendicularly, relative to the brain surface curvature of each individual recipient. Injections into mapped bV1 sites were performed using a custom-made hydraulic manipulator between ∼400-600um below the surface at 10nl/min. Recipients received ∼ 200nl of virus per injection, for a total of 2 injections. To maintain consistency, experimental agents (AAV, saline) were injected only into V1 on the right hemisphere. After injection, micropipettes were incrementally withdrawn to prevent backflow and sites resealed with Vetbond. Post-operative mice were returned to heated home cages upon waking, and received subcutaneous injections of Carprofen to reduce pain and inflammation (up to 3 days) after installation. Mice were returned to normal routine for ∼3wks while viral expression came online and stabilized.

#### Juvenile AAV

To silence Calb1 expression specifically within juvenile INs, P3-5 mice were anesthetized via hypothermia and injected with cre-dependent AAV expressing Calb1 shRNA (AAV1-CAG-DIO-eGFP-m-Calb1-shRNAmir, Vector Biolabs) into the right V1. An eGFP fluorescent tag was included in the polycistronic construct to visualize successful AAV incorporation and expression. Pup V1s were targeted using stereotaxic coordinates (3–4 mm lateral and 1 mm anterior relative to lambda) at a rate of 30–40nl/min. Due to difficulty sexing newborn mice, injected litters were randomly split between AAV recipients and saline controls. Recipient mice with off-target virus injections were excluded from subsequent experiments.

#### Adult IN transplantation and AAV

To silence Calb1 expression within INs during MGE transplantation, P120-180 VGAT-tdTomato mice were co-injected with donor MGE (prepared as described above) and cre-restricted, Calb1-overexpressing virus (AAV1-CMV-DIO-mCalb1-T2A-eGFP, UCI CNCM). After headplate installation and V1 mapping, adult VGAT-tdTomato mice were anesthetized with 2% isoflurane and secured into the surgical rig. All recipients received pre-operative Carprofen, Ringer’s, and eye ointment. Body temperature was maintained throughout the procedure using a feedback-controlled heating pad, and anesthesia gradually reduced as needed. To allow the injection micropipette to penetrate, skull underlying mapped bV1 injection sites was thinned using a dental drill (Midwest, 78044) and FG1/4 carbide burr.

31Glass micropipettes (Wiretrol 5μl, Drummond Scientific Company) were pulled with ∼75um diameter tips and beveled. Donor MGE and/or virus was front-loaded into prepared micropipettes and angled perpendicularly, relative to the brain surface curvature of each individual recipient. Injections into mapped bV1 sites were performed using a custom-made hydraulic manipulator (Narishige, MO-10) between ∼400-600um below the surface at 10nl/min. MGE only recipients were transplanted as described above. Dual recipients received ∼1 isolated MGE and 200nl of virus per injection, for a total of 2 injections. To maintain consistency, experimental agents (MGE, AAV) were injected only into V1 on the right hemisphere. After injection, micropipettes were incrementally withdrawn to prevent backflow and sites resealed with Vetbond. Post-operative mice were returned to heated home cages upon waking, and received subcutaneous injections of Carprofen to reduce pain and inflammation (up to 3 days) after installation. Mice were returned to normal routine for ∼5wks, in line with the normal timing of transplant-reactivated plasticity (35 DAT).

### Intrinsic Signal Optical Imaging (ISOI) for Ocular Dominance Plasticity

To assess whether experimental mice exhibit ocular dominance plasticity, ISOI was performed before and after monocular deprivation (MD) to measure any changes in ocular dominance. Experimenter was blinded to experimental groups from preMD imaging to post-hoc immunohistochemistry. After inducing and maintaining a steady plane of light anesthesia (0.8-1% isoflurane for adults, 0.6-0.9% isoflurane for juveniles), headplated mice were secured into the imaging rig. Headplate windows were filled with PBS and covered with a 10mm glass coverslip. PBS was added as needed throughout sessions to keep coverslips level with the top of the headplate. Eye moisture was maintained using periodic silicone oil application, and body temperature regulated by a feedback-controlled heating pad.

Intrinsic signal images were collected using a custom-designed macroscope equipped with a camera positioned over the headplate. First, a green (530 nm) LED was used to visualize and capture an image of surface vasculature. Then, the camera was refocused ∼450-550 μm below the pia surface to target cortical layer II/III of bV1, and a red (617 nm) LED used to acquire the intrinsic signal. Visual stimuli were displayed on a monitor positioned 25cm in front of the mice, aligned to their midline and eye line. To capture eye-specific responses, a single eye-block was positioned in front of one eye or the other (alternating) during stimuli presentation and recording.

Stimuli was generated using MATLAB Psychophysics Toolbox extensions and consisted of contrast-modulating sweeping noise restricted to −5° to +15° visual field azimuth and −18° to +36° visual field elevation. Each recording trial was a 5min presentation of stimuli at 0° and 180°, with a total of 3-4 recordings for each eye. Custom-written Matlab scripts using Fourier analysis were used to generate amplitude and phase maps of bV1 responses. Fourier maps were then smoothed with 5 × 5 Gaussian kernels to compute final amplitude maps. To account for hemodynamic delay, phase maps were normalized by subtracting the phase of cortical responses at 180° from the phase at 0°. Maps of absolute retinotopic phase were shown in terms of visual angle relative to the center of the monitor. Light scatter measurements were calculated from background-subtracted maps. Recordings with scatter >6 were excluded from analysis.

The ocular dominance index for each animal was calculated as (*C* − *I*)/(*C + I*), where *C* is the average response amplitude for the contralateral eye and *I* is the average response amplitude for the ipsilateral eye, each across 3-4 trials. Ocular dominance shift was calculated by subtracting PreMD ODI from PostMD indices.

### Monocular Deprivation

To reveal the presence or absence of underlying circuit plasticity, experimental mice were subjected to a temporary change in visual experience. Immediately following PreMD imaging, the eye contralateral to the injected hemisphere was sutured using Perma-Hand Silk (Ethicon, K809H). While under anesthesia, mice were quickly transferred from the imaging rig to the surgical rig. 2% isoflurane, pre-operative Carprofen, and Ringer’s were administered. Body temperature was maintained throughout the procedure using a feedback-controlled heating pad. Two mattress sutures were applied across the contralateral eyelid (one on each side) and the final knot additionally secured with a dab of Vetbond. Post-operative mice were returned to heated home cages upon waking, and received subcutaneous injections of Carprofen to reduce pain and inflammation (up to 3 days) if needed.

Eyelid sutures were monitored daily for integrity, and carefully removed after 3 days. To allow the eye to fully open before PostMD imaging, mice were returned to their heated home cage for 30min after suture removal. Mice with prematurely-opened sutures, cataracts, cloudy eyes, or drooping eyelids were excluded from PostMD imaging.

### Statistics

All quantification and data analyses were performed in a blind and unbiased manner. Statistical analyses were conducted in Prism. Experimental group numbers were determined by conservative power analysis of our previous work and other relevant studies. Datasets are first tested for normality, then appropriate statistical tests determined based on number of experimental groups and noted in individual figures.

---

**Supp. Fig 1.**
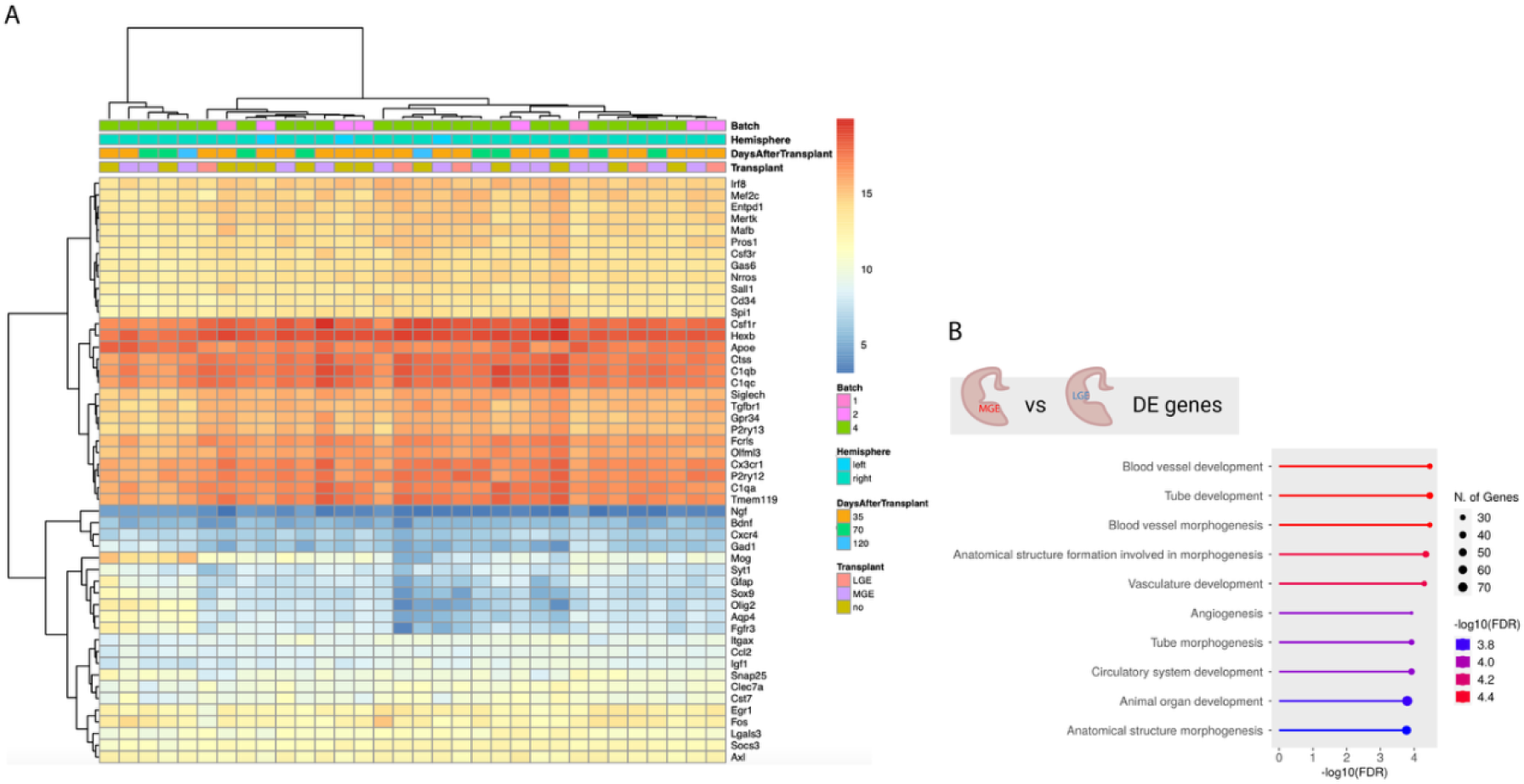
Inhibitory neuron transplantation does not activate local microglia. (A) Heatmap of select genes from microglial transcriptomes across experimental groups. Analysis conducted via DESeq2, variance stabilizing transformation applied for visualization. Selected list includes homeostatic microglial markers, inflammatory microglial markers, and aging markers (10). Negative control genes (oligodendrocyte markers, astrocyte markers) included for contrast. (B) Gene ontology annotation of enriched biological processes for the top 1% (∼300 genes by ranked p-value) of DE genes between microglia from MGE (reactivated plasticity) vs. LGE (no plasticity) recipients. Analysis conducted via ShinyGO 0.77 using the GO Biological Process database.

**Supp. Fig 2.**
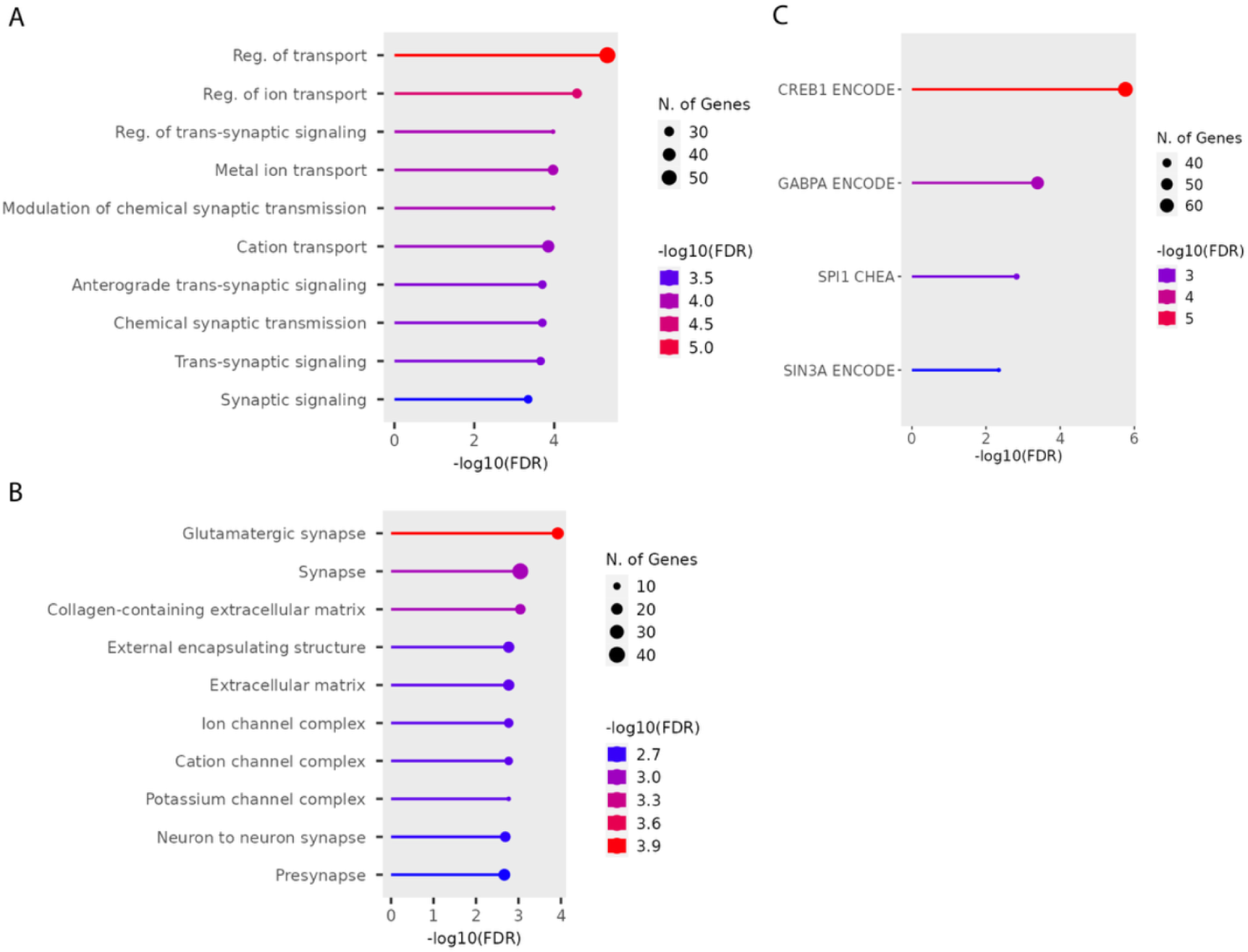
Comparison of host inhibitory neuron gene expression profiles from MGE recipients 35 DAT (peak of reactivated plasticity) vs. 70 DAT (after closure). Gene ontology annotation of enriched (A) biological processes (GO Biological Process database), (B) cellular components (GO Cellular Component database), and (C) predicted transcription factor interactions (ENCODE and ChEA chlP-X consensus database) for the top 1 % (by ranked p-value) of 35 DAT MGE vs. 70 DAT MGE DE genes (∼340). Analyses conducted via ShinyGO 0.77.

**Supp. Fig 3.**
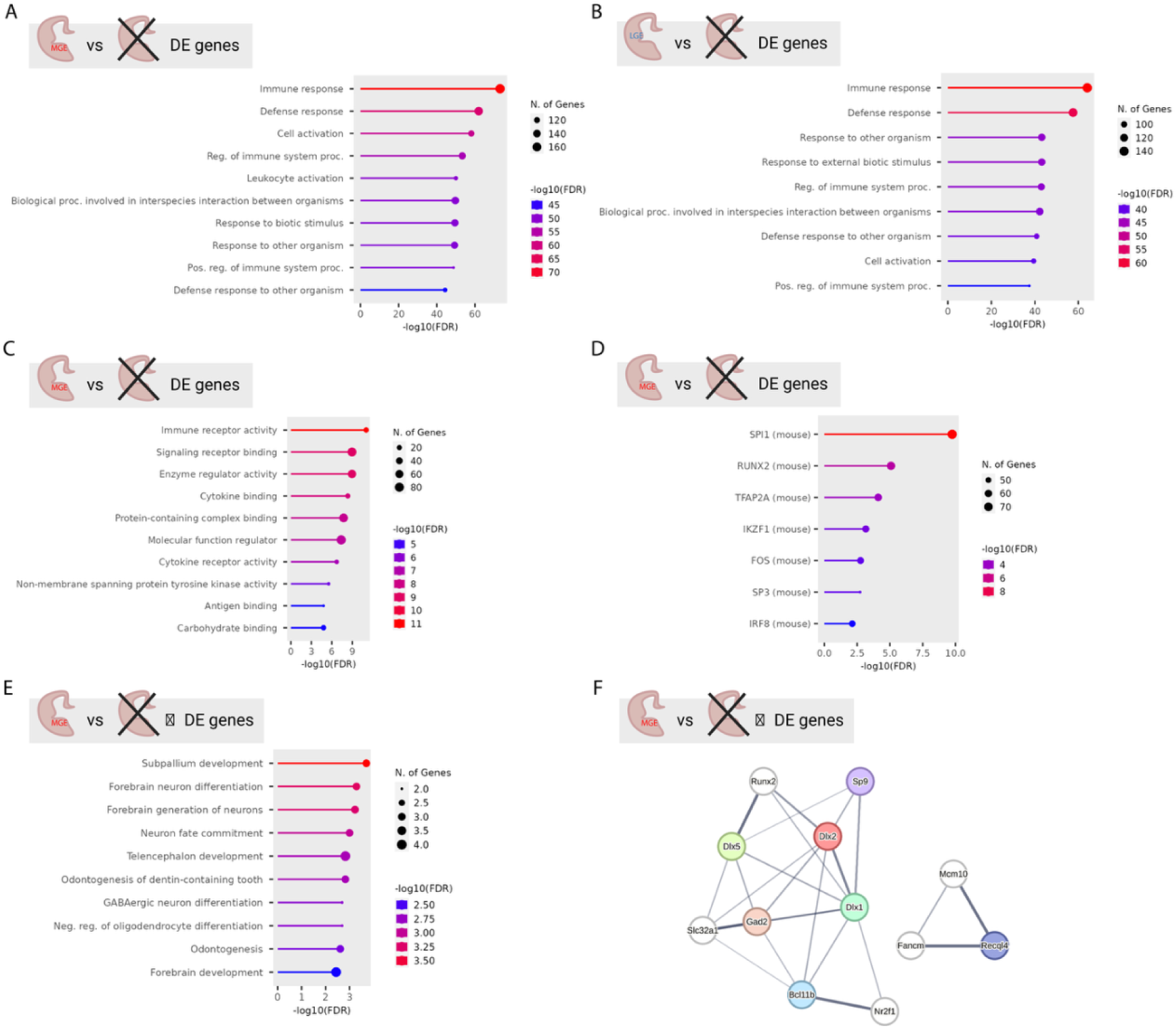
Comparison of host inhibitory neuron gene expression profiles from MGE vs. non-transplanted adult visual cortex. Gene ontology annotation of enriched biological processes for the top 1% (by ranked p-value) of DE genes between (A) 35 DAT MGE vs. 35 DAT adult no tx (∼320), and (B) 35 DAT LGE vs. 35 DAT adult no tx (∼310). Analyses conducted via ShinyGO 0.77 using the GO Biological Process database. Gene ontology annotation of enriched (C) molecular functions (GO Molecular Function database) and (D) predicted transcription factor interactions (TRANSFAC and JASPAR PWMs database) for the top 1% (by ranked p-value) of 35 DAT MGE vs. 35 DAT adult no tx DE genes (∼320). Analysis conducted via ShinyGO 0.77. (E) Gene ontology annotation of enriched molecular functions for significantly (p<0.05) downregulated genes between 35 DAT MGE vs. 35 DAT adult no tx. Analysis conducted via ShinyGO 0.77 using the GO Molecular Function database. (F) Predicted network analysis for significantly (p<0.05) downregulated genes between 35 DAT MGE vs. 35 DAT adult no tx. Analysis conducted using STRING with a minimum of med-high confidence (0.6) for interaction scores. Colored circles (nodes) represent input DE genes, lines between nodes represent functional and/or physical associations. White nodes represent the top 5 predicted interactors relative to the input network. Line weights correlate to strength of data support for predicted associations. Disconnected nodes omitted for visualization. PPI enrichment: p= 9.17e-04.

